# Asparagine starvation suppresses histone demethylation through iron depletion

**DOI:** 10.1101/2022.08.14.503884

**Authors:** Jie Jiang, Sankalp Srivastava, Sheng Liu, Gretchen Seim, Minghua Zhong, Sha Cao, Utpal Davé, Reuben Kapur, Amber L. Mosley, Chi Zhang, Jun Wan, Jing Fan, Ji Zhang

## Abstract

Nutrient availability can impact epigenome to modify gene expression and dictate cell fate decision (Etchegaray and Mostoslavsky, 2016; Kinnaird et al., 2016). α-ketoglutarate is an indispensable substrate for the Jumonji family of histone demethylases (JHDMs) mediating most of the cellular demethylation reactions on histone subunits (Schvartzman et al., 2018). Since α-ketoglutarate is an intermediate of the tricarboxylic acid (TCA) cycle and a product of transamination, its intracellular levels are regulated by the metabolism of several amino acids (Baksh et al., 2020; Carey et al., 2015; Raffel et al., 2017; Vardhana et al., 2019). Here we show that asparagine starvation suppresses global histone demethylation. This process is neither due to the change of expression of histone modifying enzymes, nor due to the change of intracellular level of α-ketoglutarate. Rather, asparagine starvation reduces intracellular pool of labile iron (Fe^2+^), which is a key cofactor for the JHDMs to function. Mechanistically, asparagine starvation post-transcriptionally suppresses the expression of iron responsive element binding protein 2 (IREB2), an iron sensing protein, which then reduces the mRNA expression of the transferrin receptor (TFRC), a major carrier for iron uptake (Hentze et al., 2010). Furthermore, iron supplementation to the culture medium restores histone demethylation and alters global gene expression to accelerate cell death under conditions of asparagine starvation. Collectively, our results uncover that suppression of iron-dependent histone demethylation is part of the cellular adaptive response to asparagine starvation.

## Introduction

Mammalian cells respond to amino acid starvation through transcriptional and translational regulation to dictate cell fate (Broer and Broer, 2017; Wek, 2018). Recent work suggests that amino acid restriction can also modulate gene expression through altering epigenetic modifications on DNA and histone, thereby impacting cell fate decision (Dai et al., 2020; Etchegaray and Mostoslavsky, 2016). For example, short-term methionine restriction poises inducible pluripotent stem cells (iPSCs) for differentiation by reducing intracellular levels of S-adenosylmethionine (SAM), which is derived from methionine and is an indispensable substrate for DNA and histone methyltransferase (Shiraki et al., 2014). Similarly, studies in multiple mammalian cell systems show that α-ketoglutarate (α-KG) is a key substrate for DNA and histone demethylases (Baksh et al., 2020; Carey et al., 2015; Klysz et al., 2015; Pan et al., 2016; Raffel et al., 2017; Vardhana et al., 2019). Since intracellular α-KG can be maintained through the metabolism of several amino acids, including glutamine, serine and branch-chained amino acids, manipulating their availability can profoundly affect cell fate decision through modulating histone and DNA demethylation.

Unlike the other proteinogenic amino acids, asparagine is the only one that has not been found to be able to be catabolized in mammalian cells (Pavlova et al., 2018), and thus whether its availability can contribute to histone methylation or demethylation has not been studied. Using human leukemic cells that are auxotrophic for asparagine as a model, we reported that asparagine starvation led to an increase of several histone methylation markers. This increase was neither due to the change of expression of histone modifying enzymes nor due to the alteration of intracellular SAM or α-KG. Instead, asparagine starvation reduces intracellular pool of labile iron (Fe^2+^), an indispensable cofactor for the JHDMs, and therefore suppresses histone demethylation. We demonstrate that asparagine starvation suppresses the expression of iron responsive element binding protein 2 (IREB2), an iron sensing protein (Pantopoulos et al., 2012), to reduce the mRNA expression of the transferrin receptor (TFRC), a major carrier for iron uptake (Hentze et al., 2010). Furthermore, iron supplementation under conditions of asparagine starvation restored histone demethylation and altered gene expression to accelerate cell death. Taken together, our results identified that regulation of iron-dependent histone demethylation is part of the metabolic adaptation during asparagine starvation.

## Results and Discussion

### Asparagine starvation leads to an increase of histone methylation

To determine the impact of asparagine availability on histone methylation status, we withdraw asparagine from the culture medium in four acute lymphoblastic leukemia (ALL) cell lines. We found that depletion of exogenous asparagine for 24-48 hours caused global increase of tri-methylation of histone H3 subunit at multiple lysine residues, including K27, K4 and K9 (Fig. 1A and Fig S1A), in RS4;11 and DND-41 cells that do not express asparagine synthetase (ASNS) (Fig. 1B). There was no change of these markers in SEMK2 and Jurkat cells that express high levels of ASNS (Fig. 1A and 1B). We also did not observe any change of di-methylation at H3K4 residue in all these four cell lines (Fig. 1A). Overexpression of ASNS in RS4;11 cells prevented the increase of H3K27me3, H3K4me3 and H3K9me2/3 following asparagine depletion (Fig. 1C); while *ASNS* deletion in Jurkat cells induced these 3 markers when exogenous asparagine was removed (Fig. 1D). These results suggest that the inability to maintain asparagine availability is the trigger of global increase of histone H3 methylation.

**Figure 1.**
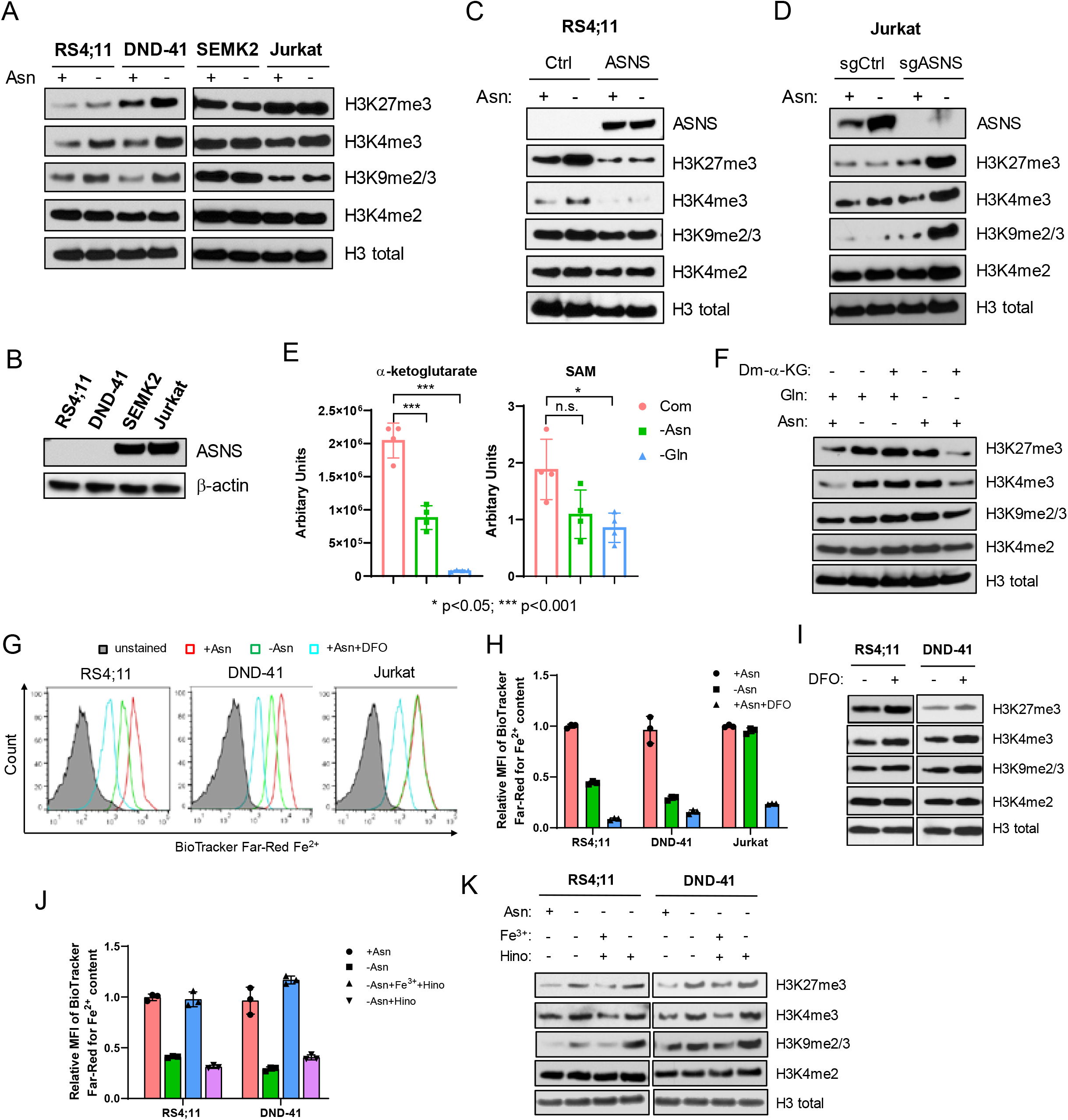
Asparagine starvation suppresses histone demethylation through iron depletion. A. RS4;11, DND-41, SEMK2 and Jurkat cells were subjected to asparagine depletion for 24 hours. Cellular expression of H3K27me3, H3K4me3, H3K9me2/3 and H3K4me2 were determined by Western Blotting. B. The expression of ASNS protein was determined by Western Blotting in RS4;11, DND-41, SEMK2 and Jurkat cells. C. ASNS was overexpressed in RS4;11 cells through lentivirus-mediated transduction. Control- or ASNS-transduced cells were subjected to asparagine depletion for 24 hours. The expression of H3K27me3, H3K4me3, H3K9me2/3 and H3K4me2 were determined by Western Blotting. D. *ASNS* gene was deleted in Jurkat by CRISPR. Control or ASNS-deleted Jurkat cells were subjected to asparagine depletion for 24 hours. Expression of H3K27me3, H3K4me3, H3K9me2/3 and H3K4me2 were determined by Western Blotting. E. RS4;11 cells were subjected to asparagine or glutamine depletion for 24 hours. Intracellular levels of α-ketoglutarate (α-KG) and S-adenosylmethionine (SAM) were determined by quantitative LC-MS. F. RS4;11 cells were subjected to asparagine or glutamine depletion for 24 hours with or without 2 mM Dimethyl-α-ketoglutarate (Dm-α-KG). Expression of H3K27me3, H3K4me3, H3K9me2/3 and H3K4me2 were determined by Western Blotting. G. RS4;11, DND-41 and Jurkat cells were subjected to the treatment for 24 hours and then stained with BioTracker Far-Red for Fe^2+^ detection by FACS. DFO was added at 100 µM in the presence of asparagine as a control. H. Quantification of the mean fluorescence intensity (MFI) of the BioTracker Far-Red staining in the panel (G). I. RS4;11 and DND-41 cells were treated with 100 µM DFO in the presence of asparagine for 24 hours. The expression of H3K27me3, H3K4me3, H3K9me2/3 and H3K4me2 were determined by Western Blotting. J. RS4;11 and DND-41 cells were subjected to asparagine depletion for 24 hours. Hinokitiol (0.75 µM) with or without Fe(NO_3_)_3_ (10 µM) was added and the MFI of BioTracker Far-Red was determined by FACS. K. RS4;11 and DND-41 cells were treated as panel (j). The expression of H3K27me3, H3K4me3, H3K9me2/3 and H3K4me2 were determined by Western Blotting. Results in panel E, H and J were shown as mean ± SD (standard derivation). p values were determined by using Student’s two-tailed unpaired t-test.

Since amino acid metabolism can affect intracellular levels of SAM and α-KG, we performed quantitative liquid chromatograph mass spectrometry (LC-MS) to measure α-KG and SAM. We found that asparagine starvation reduced α-KG to ∼40%, while glutamine starvation nearly depleted intracellular α-KG (Fig. 1E). In addition, asparagine starvation neither increased intracellular SAM or 2-hydroxylglutarate levels nor altered the ratio of α-KG/succinate or SAM/SAH (S-adenosyl-homocysteine) (Fig. 1E, Fig S1B & S1C). To determine whether the reduction of α-KG by asparagine starvation causes increased histone H3 methylation, we supplemented cells with a cell-permeable α-KG (Dm-α-KG). We found that Dm-α-KG supplementation reduced H3K27me3, H3K4me3 and H3K9me2/3 under glutamine starvation, but not under asparagine starvation (Fig. 1F), suggesting the reduction of α-KG by asparagine starvation is not the cause of increased histone H3 methylation. In addition, we found no correlative changes in the expression of histone methyltransferases or demethylases that could consistently explain the change of these 3 markers during asparagine starvation (Fig S1D).

### Asparagine starvation reduces intracellular pool of labile iron (Fe^2+^) and thereby suppresses histone demethylation

Since the demethylation of H3K27me3, H3K4me3 and H3K9me2/3 requires JHDMs that also use Fe^2+^ as a co-factor, we measured intracellular Fe^2+^ levels using BioTracker Far-red, a dye specific for ferrous (Hirayama et al., 2017). We used deferoxamine (DFO), an iron chelator, to treat the cells in complete medium as a negative control. We found that asparagine starvation for 24 hours reduced Fe^2+^ content to less than 50% in RS4;11 and DND-41 cells, but not in Jurkat cells (Fig. 1G and 1H). Consistent with the literature (Rensvold et al., 2016), DFO treatment induced H3K27me3, H3K4me3 and H3K9me2/3 in RS4;11 and DND-41 cells, with no impact on H3K4me2 (Fig. 1I). Since demethylation of H3K4me2 uses LSD1/2, two lysine specific demethylases that do not use iron as a co-factor (Shi and Whetstine, 2007), our results are consistent with the hypothesis that the global increase of histone H3 methylation under asparagine starvation is due to a defect of iron-dependent histone demethylation. To determine whether the reduction of intracellular Fe^2+^ is necessary for the increase of histone H3 methylation, we treated RS4;11 and DND-41 cells with Fe^3+^ in the presence of Hinokitiol, a chemical carrier for iron import (Grillo et al., 2017). We found that Fe^3+^ plus Hinokitiol, but not Hinokitiol alone, restored intracellular Fe^2+^ content and histone H3 demethylation (Fig. 1J and 1K).

### Asparagine starvation reduces the cell surface expression of the transferrin receptor (TFRC)

To determine the cause of the reduction of intracellular Fe^2+^ pool, we analyzed published RNA-seq data from RS4;11 cells subjected to asparagine starvation (Jiang et al., 2019) and found reduced expression of iron homeostatic genes (Fig. 2A and 2B). We confirmed the reduction of the mRNAs of *TFRC, STEAP3, SFXN2, SLC46A1* (Fig. S2A), whose major roles are iron uptake and intracellular transport. Consistently, both total cellular expression and cell surface expression of transferrin receptor (TFRC), also known as CD71, was reduced by asparagine starvation in RS4;11 and DND-41 cells, but not in SEMK2 and Jurkat cells (Fig. 2C-2E). Furthermore, ASNS overexpression restored TFRC cell surface expression in RS4;11 cells when asparagine was withdrawn; while *ASNS* deletion in Jurkat cells reduced TFRC cell surface expression under the same condition (Fig. S2B & S2C). Since TFRC is a major carrier for iron uptake (Hentze et al., 2010), we hypothesize that the reduction of TFRC expression by asparagine starvation leads to the reduction of intracellular Fe^2+^ to suppress iron-dependent histone demethylation. To test this hypothesis, we used doxycycline-inducible shRNA to suppress TFRC expression (Fig. 2F, and Fig. S2D) and found that doxycycline treatment induced H3K27me3 and H3K4me3 in the presence of exogenous asparagine (Fig. 2G). This induction was mitigated by iron supplementation in the presence of Hinokitiol (Fig. 2G), which correlated with intracellular Fe^2+^ content (Fig. S2E). Conversely, TFRC overexpression in RS4;11 and DND-41 cells restored cell surface expression of TFRC, intracellular Fe^2+^ content and partially rescued histone demethylation under asparagine depletion (Fig. S2F & S2G, and Fig 2H). These results suggest that TFRC is both necessary and sufficient for iron-dependent histone demethylation.

**Figure 2.**
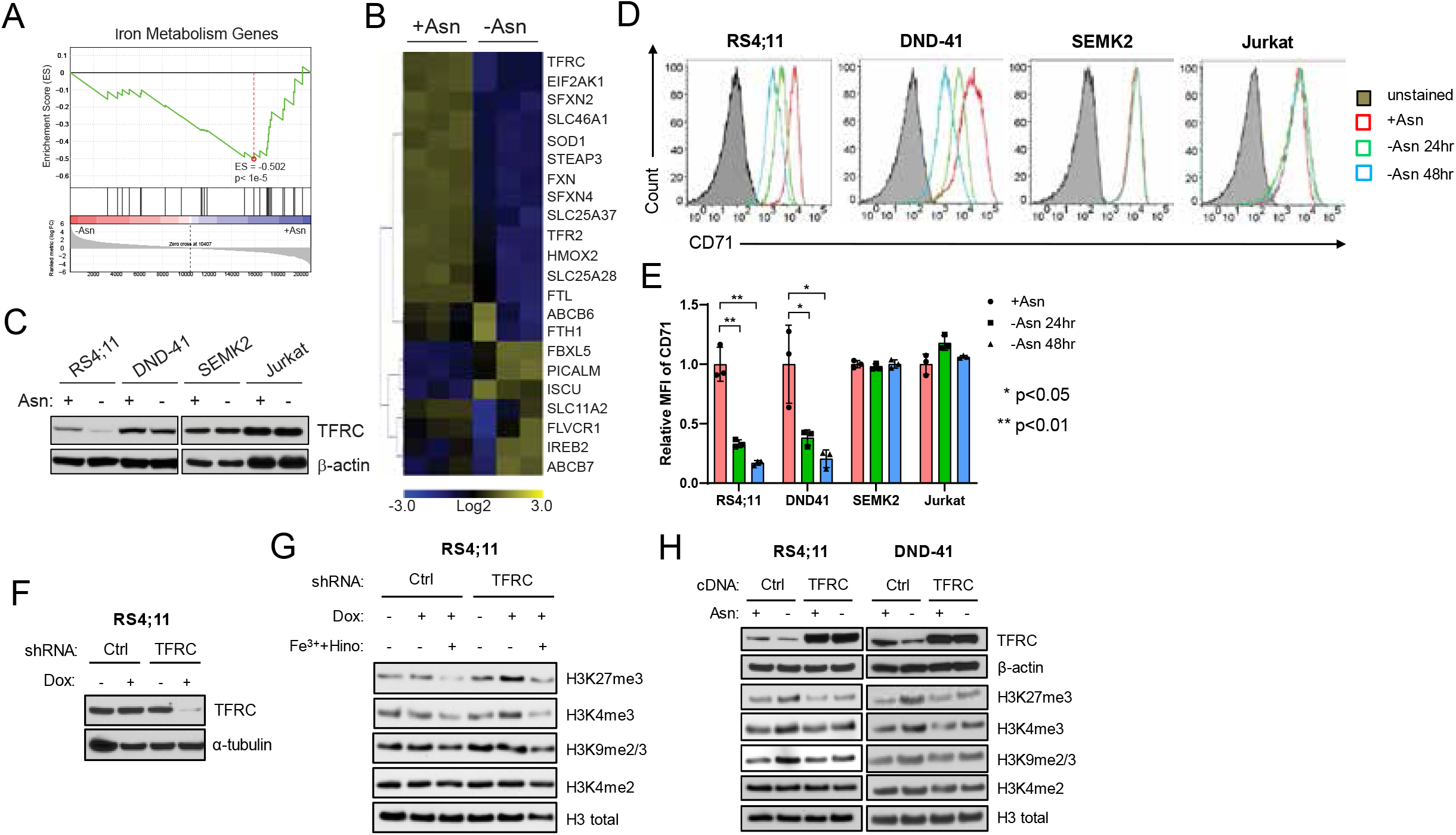
Asparagine starvation reduces intracellular iron content through downregulating the transferrin receptor (TFRC/CD71). A. Geneset Enrichment Analysis identified downregulation of mRNAs of iron metabolism genes in RS4;11 cells following asparagine depletion for 24 hours. B. Heatmap illustration of the change of expression of iron metabolism genes in panel (A). C. RS4;11, DND-41, SEMK2 and Jurkat cells were subjected to asparagine depletion for 24 hours. TFRC protein expression was determined by Western Blotting. D. RS4;11, DND-41, SEMK2 and Jurkat cells were subjected to asparagine depletion for 24 hours. Cell surface expression of TFRC/CD71 was determined by FACS analysis. E. The MFI of cell surface staining of TFRC/CD71 in panel (D) was quantified. Results were shown as mean ± SD (standard derivation). p values were determined by using Student’s two-tailed unpaired t-test. F. The expression of TFRC in RS4;11 cells was suppressed by doxycycline-inducible shRNA following 48 hours of doxycycline (1 µg/mL) treatment. G. RS4;11 cells with control or TFRC shRNA were treated with doxycycline for 48 hours with or without Hinokitiol (0.75 µM) plus Fe(NO_3_)_3_ (10 µM). The expression of H3K27me3, H3K4me3, H3K9me2/3 and H3K4me2 were determined by Western Blotting. H. TFRC cDNA was introduced into RS4;11 and DND-41 via lentivirus-mediated gene transduction. Control- or TFRC-transduced cells were subjected to asparagine depletion for 24 hours. The expression of H3K27me3, H3K4me3, H3K9me2/3 and H3K4me2 were determined by Western Blotting.

### Downregulation of IREB2 is responsible for the reduction of TFRC expression following asparagine depletion

The stability of *TFRC* mRNA is tightly regulated by iron sensing proteins. ACO1 and IREB2 are the two known iron sensing proteins that bind to the iron-responsive elements (IREs) within the 3’ UTR of *TFRC* mRNA to increase its stability when intracellular iron content is low (Pantopoulos et al., 2012). Elevated cellular iron inactivates ACO1 and IREB2 via different molecular machineries to cause their dissociation from *TFRC* mRNA, leading to its degradation and therefore providing a feedback mechanism to maintain iron homeostasis. We found that, unlike *TFRC*, the mRNA levels of *ACO1* and *IREB2* did not change following asparagine depletion in both RS4;11 and DND-41 cells (Fig. 3A). However, the protein expression of IREB2 was consistently reduced by asparagine starvation in RS4;11 and DND-41 cells, but not in SEMK2 and Jurkat cells (Fig. 3B). Similarly, only the IREB2 protein, but not ACO1, was reduced by asparagine starvation in Jurkat cells lacking *ASNS* gene (Fig. 3C). Using the doxycycline-inducible shRNAs, we found that only suppression of IREB2 reduced the expression of TFRC at both mRNA and protein levels (Fig S3, Fig 3D & 3E).

**Figure 3.**
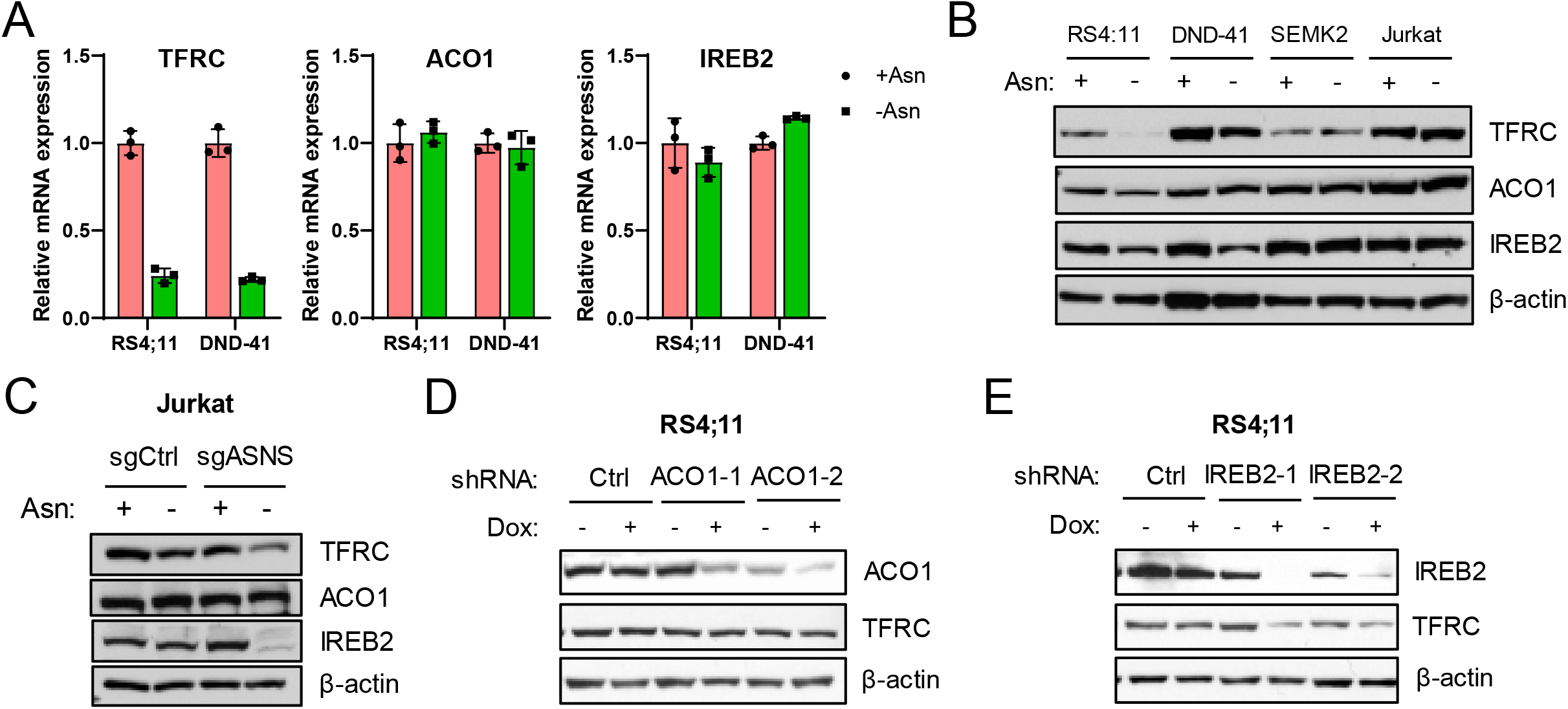
Asparagine starvation downregulates the expression of IREB2 post-transcriptionally. A. The mRNA expression of TFRC, ACO1 and IREB2 was determine by qPCR in RS4;11 and DND-41 cells following 24-hour asparagine depletion. B. RS4;11, DND-41, SEMK2 and Jurkat cells were subjected to asparagine depletion for 24 hours. The expression of TFRC, ACO1 and IREB2 protein was determined by Western Blotting. C. Control or *ASNS*-deleted Jurkat cells were subjected to asparagine depletion for 24 hours. The expression of TFRC, ACO1 and IREB2 protein was determined by Western Blotting. D. RS4;11 cells were transduced with two inducible hairpins of ACO1 and then treated with doxycycline (1 µg/mL) for 48 hours. The expression of ACO1 and TFRC protein was determined by Western Blotting. E. RS4;11 cells were transduced with two inducible hairpins of IREB2 and then treated with doxycycline (1 µg/mL) for 48 hours. The expression of IREB2 and TFRC protein was determined by Western Blotting.

### Asparagine starvation increase H3K4me3 deposition to alter gene expression and dictate cell fate

Since histone methylation profoundly affects gene expression, we hypothesize that increased H3 lysine methylation under asparagine starvation modulates gene expression to mitigate the stress. Using H3K4me3 ChIP-seq, we found a global increase of its deposition across the genome following asparagine depletion (Fig S4A). The enrichment of H3K4me3 on gene promoters was positively correlated with gene expression, consistent with the known role of H3K4me3 on transcriptional activation (Fig S4B). Furthermore, supplementation of Fe^3+^ in the presence of Hinokitiol reduced the H3K4me3 deposition in the promoter regions of a portion of genes whose H3K4me3 deposition increased in the promoter regions under asparagine starvation (Fig S4C). There was a weak correlation between the reduction of H3K4me3 deposition and gene expression alteration (Fig S4C).

The genes with increased mRNA expression or increased H3K4me3 deposition upon asparagine starvation were significantly enriched in cell periphery, plasma membrane synapse, and vesicle trafficking given either ChIP-seq (Fig. 4A) or RNA-seq results (Fig. 4B), indicating a potential enhancement of exchanging with extracellular environment. For example, genes in the cytoplasmic vesicle membrane pathway were induced by asparagine starvation (Fig. 4C). The functions of these genes are broadly linked to membrane internalization, endocytosis and endosomal function, which can facilitate the utilization of extracellular protein as a source of amino acid to mitigate the stress (Palm and Thompson, 2017). In addition, Fe^3+^ supplementation suppressed the induction of some of these genes involved in cell surface receptor recycling or vesicle trafficking, such as SLA and FGD2 (Dragone et al., 2006; Huber et al., 2008) (Fig. 4D, left panel). Furthermore, the mRNA expression of SLA and FGD2 positively correlated with H3K4me3 deposition in their promoter regions under these three conditions (Fig. 4D, right panel)

**Figure 4.**
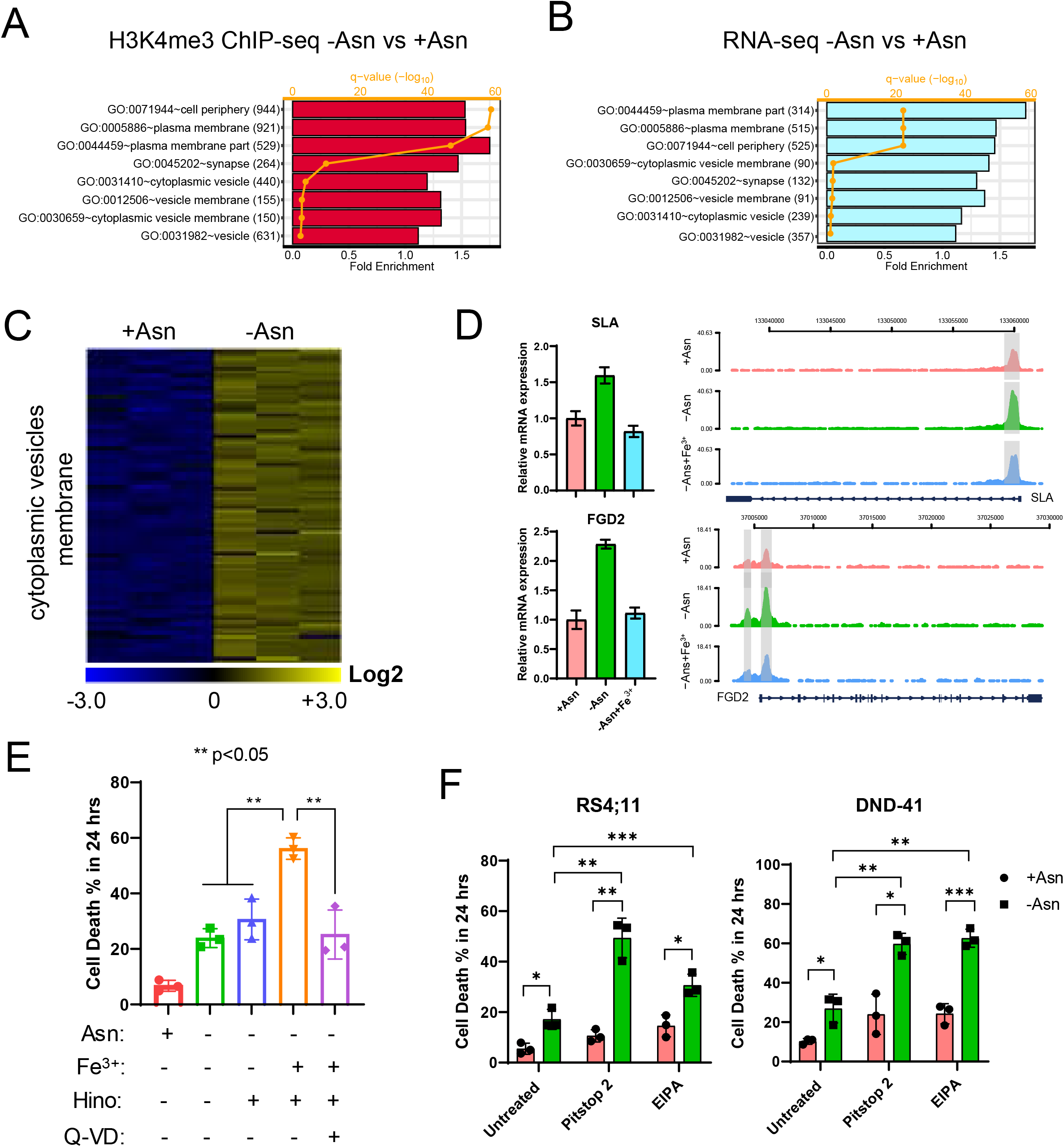
Asparagine starvation leads to an increase of H3K4me3 deposition to modulate gene expression. A. RS4;11 cells were starved for asparagine for 24 hours. H3K4me3 ChIP-seq analysis was performed (n=3). The differential expression peaks (FDR<0.05, |log2FC|>0.25) were subjected to gene ontology and pathway functional analysis by using DAVID. B. RS4;11 cells were starved for asparagine for 24 hours. RNA-seq analysis was performed (n=3). The differential expression genes (FDR<0.05, |log2FC|>0.25) were subjected to gene ontology and pathway functional analysis by using DAVID. C. RS4;11 cells were starved for asparagine for 24 hours. RNA-seq analysis was performed (n=3). Heatmap of the genes in the cytoplasmic vesicle membrane pathway was shown. D. RS4;11 cells were subjected to asparagine starvation with or without Fe^3+^ supplementation for 24 hours. RNA was harvested and the relative expression of *SLA* and *FGD2* were determined by qPCR (left). H3K4me3 ChIP-seq analysis was performed under the same conditions and relative enrichment of H3K4me3 around gene promoters was illustrated by genome browser (right). The result is a merged peak of 3 replicates. E. RS4;11 cells were subjected to asparagine starvation for 24 hours in the presence of other treatment as shown in the legend. Cell death index was measured by trypan blue staining. F. RS4;11 and DND-41 cells were cultured in asparagine-replete or -deficient media for 24 hours. Pitstop (50 µM) or EIPA (25 µM) was added in both conditions. Cell death index was measured by trypan blue staining. Results in panel E and F were shown as mean ± SD (standard derivation). p values were determined by using Student’s two-tailed unpaired t-test.

We hypothesize that cells may uptake extracellular proteins as a means to regenerate intracellular amino acid to mitigate asparagine starvation. Consistent with this hypothesis, Fe^3+^ supplementation enhanced cell death within 24 hours following asparagine withdrawal (Fig. 4E). The enhanced cell death is not due to ferroptosis (Stockwell et al., 2017), as we did not observe increased reactive oxygen species (ROS) (Fig. S4D). Furthermore, Q-VD, a pan-caspase inhibitor, suppressed the enhanced cell death triggered by Fe^3+^ supplementation (Fig. 4E). To examine our hypothesis, we treated RS4;11 and DND-41 cells with inhibitors of endocytosis (Pitstop 2) or macropinocytosis (EIPA). The results clearly showed an enhanced cell death under asparagine starvation (Fig. 4F).

We conclude that asparagine starvation in asparagine auxotrophic cells inhibits the expression of TFRC and thus suppresses iron uptake and iron-dependent histone demethylation. Iron supplementation can restore histone demethylation and therefore alter gene expression under asparagine starvation. Our results indicate that inhibition of histone demethylation is part of the metabolic adaptive response to ensure proper gene expression to mitigate the stress, as iron supplementation accelerates cell death under asparagine starvation. We identified that downregulation of IREB2 is a key molecular component to be responsible for asparagine starvation-induced inhibition of TFRC expression. However, the mechanism underlying the post-transcriptional inhibition of IREB2 expression by asparagine starvation remains to be determined. Since the connection between amino acid restriction and epigenetic regulation only comes to the focus recently (Kinnaird et al., 2016; Schvartzman et al., 2018), our study will provide a comprehensive understanding of nutrient availability and histone modifications. Furthermore, the dynamic change of TFRC/CD71 expression in a variety of physiological and pathological conditions may indicate a requirement of iron-dependent histone demethylation to regulate epigenome (Crielaard et al., 2017; Waickman and Powell, 2012). Therefore, our results uncover that iron homeostasis is another layer of control between amino acid restriction, epigenome and cell fate decision.

## Materials and Methods

We will share all research materials that are published in this study. RNA-seq and ChIP-seq data will be shared using the NCBI GEO Web Deposit Guide. Reagents, including but are not limited to cell lines, plasmid constructs, and protocols for newly developed assays, will be made available to the academic community, if necessary, through appropriate Material Transfer Agreements with terms similar to those prescribed by NIH guidelines.

## Key Resources Table

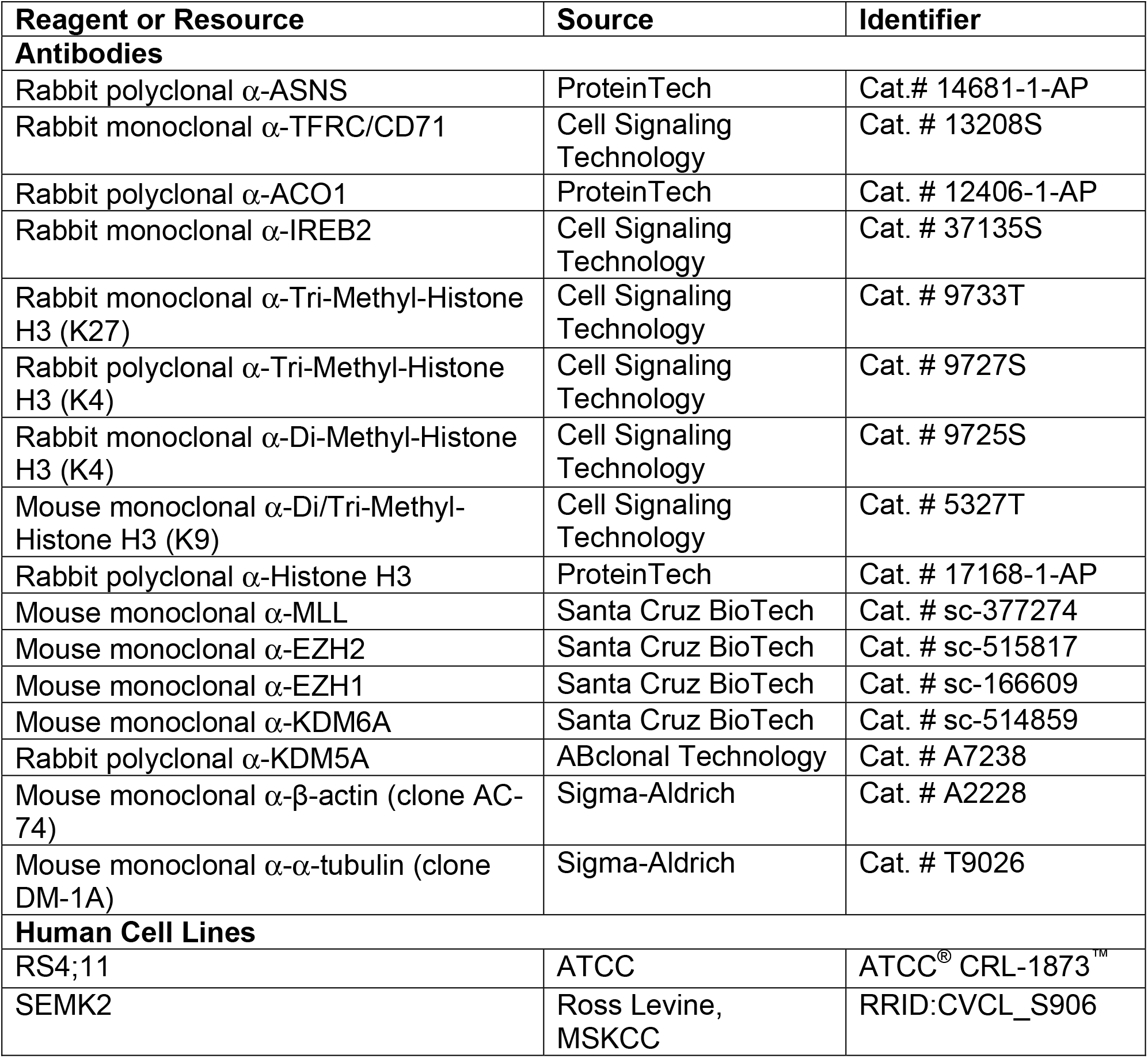
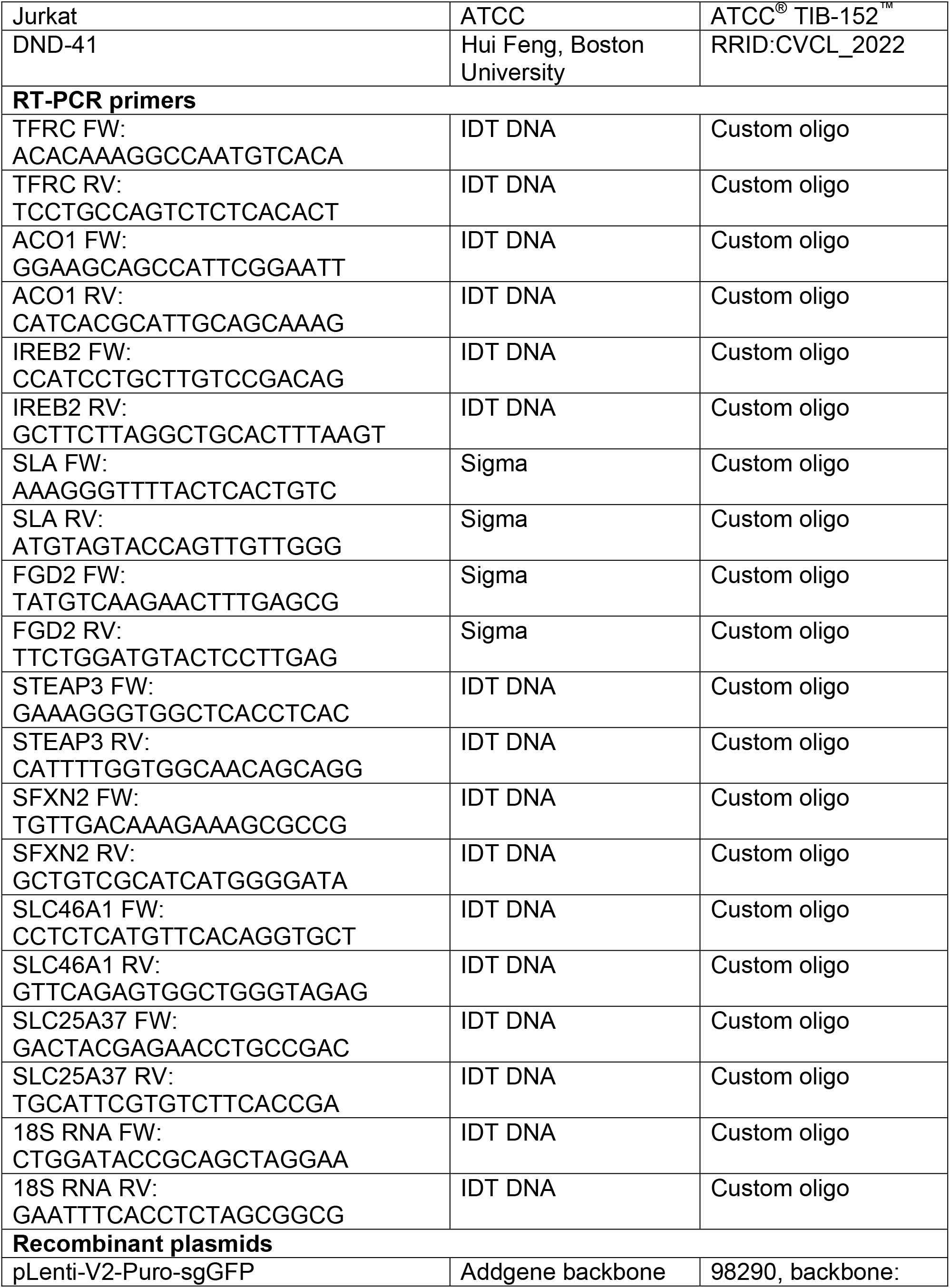
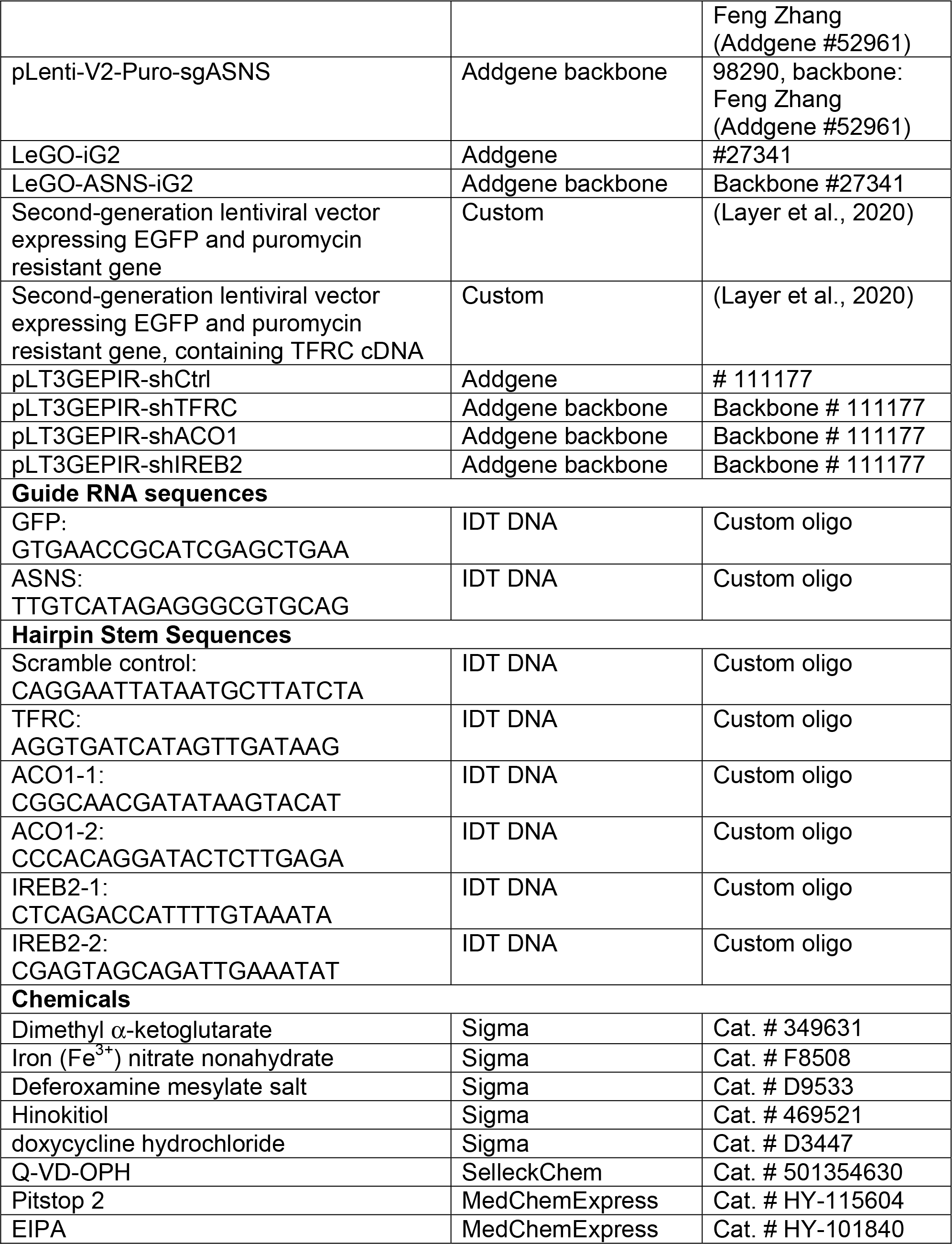

### Experimental Model and Subject Details

#### Cell Culture

All human leukemic cell lines were cultured at 37°C in 5% CO_2_ in a lymphocyte culturing medium (LCM). The base of LCM is a 1:1 ratio mixture of DMEM and IMDM. For practical reason and quality control purpose, we can consistently generate LCM in lab through supplementing high glucose DMEM (11965092, Thermo Fisher Scientific) with: 0.14 mM L-Alanine, 0.17 mM L-Proline, 5.3 × 10^−5^ mM Biotin, 9.59 × 10^−5^ mM Vitamin B12 and 9.8 × 10^−5^ mM Sodium Selenite, 10 mM HEPES, and 55 µM β-mercaptoethanol, 100 Units/mL penicillin/streptomycin, 2 mM L-Glutamine, 10% FBS and 0.1mM L-Asparagine. For asparagine or glutamine starvation, cells were centrifuged and supernatant was removed by aspiration. Cell pellets were resuspended in asparagine- or glutamine-free LCM. Asparagine- or glutamine-free LCM was made from high glucose DMEM lacking glutamine (11960044, Thermo Fisher Scientific) as described above, with the exception of using 10% dialyzed FBS. Cell viability and viable cell numbers were recorded in using the Vi-CellXR cell viability analyzer (Beckman Coulter).

#### Western Blotting

For non-histone blots, protein was extracted by using 1× RIPA buffer (diluted from 10x RIPA lysis buffer, Millipore, Cat. # 20-188) with protease inhibitors (Thermo Scientific, Cat. # 1860932) and phosphatase inhibitors (Thermo Scientific, Cat. # 78428). For histone blots, cells were first lysed with Triton Extraction Buffer (PBS containing 0.5% Triton plus protease inhibitors). Nuclei were collected by centrifugation and then lysed with 0.2 N HCl overnight at 4°C. Supernatant was collected by centrifugation and then neutralized with 1/10 volume of 2 M NaOH. Total proteins of equal amount (20 µg for non-histone and 5 µg for histone) were separated on NuPAGE Bis-Tris gels (Invitrogen, Cat. # NP0322BOX) and then transferred to Nitrocellulose membranes (Bio-Rad, Cat. # 1620115). Membranes were blocked in 5% milk and then incubated with corresponding primary antibodies overnight at 4°C. Membranes were washed with 1×Tris Buffered Saline with Tween 20/TBST (diluted from 20x TBST, Santa Cruz Biotechnology, Cat. # 362311) and then incubated with horseradish peroxidase (HRP) conjugated secondary antibody (ECL anti-rabbit IgG, Sigma, Cat. # NA934V; ECL anti-mouse IgG, Sigma, Cat. # NA931V. 1:5000 dilution). Membranes were washed with 1× TBST and subjected to Chemiluminescent Western ECL detection (Thermo Scientific, Cat. # 32106). The blots were stripped with Restore Western Blot Stripping Buffer (Thermo Scientific, Cat. # 21059), washed with 1xTBST, and then re-probed with appropriate primary antibodies for signal detection.

#### Mass Spectrum Analysis of Intracellular Metabolite

Cells were collected by centrifugation, supernatant was aspirated and the pellets were washed thoroughly once with ice cold 1 × HBSS (14025092, Life Technology). Cellular metabolites were extracted with 80% methanol on ice. Supernatant was collected and dried with SpeedVac (SPD111V, Thermo Fisher Scientific) connected to Refrigerated Vapor Trap (RVT5105, Thermo Fisher Scientific) at room temperature. Dried samples were resuspended in 40:40:20 acetonitrile:MeOH:water and analyzed using a Thermo Q-Exactive mass spectrometer coupled to a Vanquish Horizon UHPLC. Metabolites were separated on a 150 × 2.1 mm Xbridge BEH Amide (2.5µM) HPLC Column (Waters). Samples were run with a gradient of solvent A (95% H_2_O, 5% ACN, 20mM NH_4_AC, 20mM NH_4_OH) and solvent B (20% H_2_O, 80% ACN, 20mM NH_4_AC, 20mM NH_4_OH) as follows: 0 min, 100% B; 3 min, 100% B; 3.2 min, 90% B; 6.2 min, 80% B; 10.5 min, 80% B, 10.7 min, 70% B; 13.5 min, 70% B; 13.7 min, 45% B; 16 min, 45% B; 16.5 min 100% B; 22 min, 100% B. Data were collected on a full scan. Flow rate was 0.2 ml/min. Metabolites were identified based on exact M/z and retention time determined using chemical standards. Data were analyzed with Maven (Clasquin et al., 2012; Melamud et al., 2010), and normalized to internal standard of ^13^C_4_,^15^N_2_-Asparagine (1 pmole/sample) and then total cell number of each sample.

#### Intracellular Ferrous (Fe^2+^) and ROS Measurement

Cells were collected by centrifugation and washed once with PBS. BioTracker Far-Red (SCT037, Millipore Sigma) was diluted in serum-free DMEM at final concentration of 5 µM. Resuspend 1 million cells in 100 µL of diluted BioTracker Far-Red staining medium in 96-well plates and incubate the plate reaction in 37°C CO_2_ incubator for 1 hour. Cells with staining medium was further diluted with 400 µL serum-free DMEM containing eFluor 450 (65-0863-14, eBioscience) at 1:4000 dilution for dead cell labeling. The signal of BioTracker Far-Red was captured in RL1 channel by the Invitrogen Attune NxT Flow Cytometer. For the ROS measurement, cells were stained in 100 µL serum-free DMEM with 5 µM H_2_DCFDA (D399, Thermo Fisher Scientific) at 37°C for 30 minutes and then diluted with 400 µL serum-free DMEM before detection. The signal of H_2_DCFDA was captured in BL1 channel by the Invitrogen Attune NxT Flow Cytometer. We used APC anti-human CD71 (334108, BioLegend Inc.) to label cell surface expression of CD71.

#### Molecular Cloning and Virus Production

Mouse *ASNS* cDNA was cloned into the LeGO-iG2 vector backbone for overexpression in RS4;11 cells (Addgene, #27341). Human TFRC/CD71 cDNA was cloned into a second-generation lentiviral vector expressing EGFP and puromycin resistant gene at the same time for dual-selection (Layer et al., 2020). Guide RNAs for human *ASNS* gene were designed using Feng Zhang lab’s CRISPR design resource: http://crispor.tefor.net/ and cloned into pLentiCRISPRv2-Puro vector (Addgene, Cat. # 98290). The design of the doxycycline-inducible shRNAs targeting human TFRC, ACO1 and IREB2 was previously described (Pelossof et al., 2017). The hairpins were cloned into the LT3GEPIR vector (Addgene, Cat# 111177) (Fellmann et al., 2013). We used pMD2.G (Addgene, Cat# 12259) and psPAX2 (Addgene, Cat# 12260) as packaging plasmids for lentiviral particle production in 293T cells.

#### mRNA Quantification

Total RNA was then isolated with TRIzol (15596026, Life Technologies) according to the manufacturer’s instructions. 0.5∼2 µg total RNA was processed for cDNA synthesis with random hexamer primers, using EasyScript Plus RTase from the EasyScript Plus cDNA Synthesis Kit (Lamda Biotech, Cat. # G235). The synthesized cDNA was then subjected for qPCR amplification with designed PCR primers for human target genes.

#### RNA Sequencing

Total RNA was first evaluated for its quantity, and quality, using Agilent Bioanalyzer 2100. For RNA quality, a RIN number of 7 or higher is desired. One hundred nanograms of total RNA was used. cDNA library preparation included mRNA purification/enrichment, RNA fragmentation, cDNA synthesis, ligation of index adaptors, and amplification, following the KAPA mRNA Hyper Prep Kit Technical Data Sheet, KR1352 – v4.17 (Roche Corporate). Each resulting indexed library was quantified and its quality accessed by Qubit and Agilent Bioanalyzer, and multiple libraries pooled in equal molarity. The pooled libraries were then denatured, and neutralized, before loading to NovaSeq 6000 sequencer at 300pM final concentration for 100b paired-end sequencing (Illumina, Inc.). Approximately 30-40M reads per library was generated. A Phred quality score (Q score) was used to measure the quality of sequencing. More than 90% of the sequencing reads reached Q30 (99.9% base call accuracy).

#### Chromatin Immunoprecipitation Followed by High Throughput Sequencing (ChIP-seq)

1 × 10^7^ cells were resuspended in 25 mL PBS and fixed with 1% formaldehyde (final concentration) on a platform rocker at room temperature for 10 minutes. 1.4mL of 2.5M glycine was added and incubated for another 5 minutes on platform rocker to quench the crosslinking reaction. Cells were washed and lysed in 2mL cell lysis buffer A (20 mM Tris-HCl pH=8.0, 85 mM KCl, and 0.5% NP-40) and incubated on ice for 10 minutes. Nuclei was pelleted by centrifugation at 1,350 g for 5 minutes at 4 °C. The nuclei was then resuspended in 750μl lysis buffer B (50 mM Tris-HCl pH=8.0, 10 mM EDTA, 1% SDS, plus protease inhibitor cocktail) and sonicated by using the Covaris S2 Focused-ultrasonicator until majority of DNA fragments were between 200 and 500 base pairs in size. The sonicated materials were then centrifuged at 20,000 g for 10 minutes at 4°C to collect supernatant. 10% of the supernatant (75 μL) was used as input and 300 μL supernatant was used for each immunoprecipitation (IP) reaction.

The 300 μL supernatant sample was diluted 5-fold by adding 1.2ml IP dilution buffer (1.25% Triton X-100, 187.5 mM NaCl, 20 mM Tris-HCl pH=8.0, plus protease inhibitor cocktail), and incubated overnight at 4°C by rotating with the 30 μL protein G Dynabeads (Life Technologies) pre-incubated with H3K4me3 antibody (Active Motif, 39016). The beads were washed consecutively with 1 mL Low-salt wash buffer (0.1% SDS, 1% Triton X-100, 2 mM EDTA, 20 mM Tris-HCl pH 8.0, and 150 mM NaCl, twice), High-salt wash buffer (0.1% SDS, 1% Triton X-100, 2 mM EDTA, 20 mM Tris-HCl pH 8.0, 500 mM NaCl, once), LiCl wash buffer (0.25 M LiCl, 1% NP-40, 1% Sodium Deoxycholate, 1 mM EDTA, 10 mM Tris-HCl pH 8.0, once) and TE wash buffer (50mM NaCl, 10 mM Tris pH 8.0, 1 mM EDTA, once) and incubated with 125 µL Elution Buffer (1% SDS/0.1 M sodium bicarbonate) for 15min on thermomixer at 1000 RPM and 65°C. Supernatant was collected by magnet separation from the beads. 5μL 5M NaCl was added to each elute and incubated at 65°C overnight. 30 μL of input samples were incubated with 95 μL Elution Buffer and 5 μl 5M NaCl was used for reverse crosslinking. 2µl of RNase A (0.5 mg/mL) was added to each IP and input sample for 30 minutes incubation at 37°C. 2 µL of Proteinase K (20 mg/mL) was then added to each sample and incubated for 2 hours at 55°C. PCR Purification Kit (Qiagen) was used to recover DNA from each sample in 40 μL EB buffer.

Traces of eluted DNA were used to library preparation using Illumina TruSeq Nano DNA LT Library Prep Kit (Cat# FC-121-4001), including end-repair, dA-tailing, indexed adaptor ligation and amplification. Each resulting indexed library were quantified and its quality accessed by Qubit and Agilent Bioanalyzer, and multiple libraries were pooled in equal molarity. The pooled libraries were then denatured, and neutralized, before loading onto NovaSeq 6000 sequencer at 300pM final concentration for 100b paired-end sequencing (Illumina, Inc.). Approximately 15-20M reads per library was generated. A Phred quality score (Q score) was used to measure the quality of sequencing. More than 90% of the sequencing reads reached Q30 (99.9% base call accuracy).

#### RNA-seq and ChIP-seq Analysis and Pathway Enrichment Analysis

RNA-seq: The reads were mapped to the human genome hg38 using STAR (v2.7.2a) (Dobin et al., 2013). RNA-seq aligner with the following parameter: “--outSAMmapqUnique 60”. Uniquely mapped sequencing reads were assigned to Gencode 31 gene using featureCounts (v1.6.2) (Liao et al., 2014) with the following parameters: “-s 2 –p –Q 10 -O”. The data was filtered using read count > 10 in at least 3 of the samples, normalized using TMM (trimmed mean of M values) method and subjected to differential expression analysis using edgeR (v3.36.0) (McCarthy et al., 2012; Robinson et al., 2010). Gene ontology and pathway functional analysis was performed on differential expression gene with false discovery rate cut-off of 0.05 and absolute value of log2 of fold change cut-off of 0.25 using DAVID (Dennis et al., 2003; Huang da et al., 2009). Gene set enrichment analysis was conducted by using hypergeometric test against human gene ontology and MsigDB v.6 canonical pathways, with p < 0.01 as the significant cutoff (Subramanian et al., 2005).

ChIP-Seq: Bowtie2(Langmead and Salzberg, 2012) was used for ChIP-Seq reads alignment on the human genome (hg38). Duplicated reads were removed using Picard [Ref 1]. Peak calling of mapped ChIP-Seq reads were performed by MACS2 (Zhang et al., 2008) compared with input ChIP-Seq with a bonferroni adjusted cutoff of p value less than 0.01. Peaks called from multiple samples were merged. Merged peaks overlapping with ENCODE blacklist regions (Amemiya et al., 2019; Consortium, 2012) were removed to form a final set of regions. Reads overlapping with these regions in different samples were counted by featureCounts (Liao et al., 2014). The data was filtered using at least 10 read counts in more than one of the samples, normalized using TMM (trimmed mean of M values) method and subjected to differential analysis using edgeR (v3.36.0) (McCarthy et al., 2012; Robinson et al., 2010). Gene ontology and pathway functional analysis was performed on genes whose upstream 2kbp having differential peak signals with false discovery rate cut-off of 0.05 and absolute value of log2 of fold change cut-off of 0.25 using DAVID (Dennis et al., 2003).

Ref. 1: Broad Institute. (Accessed: 2018/02/21; version 2.17.8). “Picard Tools.” Broad Institute, GitHub repository. http://broadinstitute.github.io/picard/.

#### Data rigor and reproducibility

Each experiment was performed in at least three technical replicates and the experiment was repeated at least once at a different time as biological replicates to confirm observed phenotypes and data reproducibility. All western blots shown were repeated in at least two independent experiments, with a representative blot shown in the figures. GraphPad Prism (v9.2.0) was used to plot data and perform statistical analysis as described.

## Acknowledgement

JZ is supported by NIH/NCI CA244625, Showalter Trust Fund and Riley Children Foundation. There are no competing interests to report. We thank the IUSCC Flow Cytometry Core for the cell sorting. We thank the IU Simon Cancer Center (Grant P30CA082709), Purdue University Center for Cancer Research (Grant P30CA023168) and Walther Cancer Foundation to support our data analysis through the Collaborative Core for Cancer Bioinformatics (C^3^B).

## Author Contributions

JJ, JZ wrote the manuscript. JJ and SS designed and performed experiments, and analyzed the results. SL, GS, MZ, SC performed experiments. UD, RK and AM provided critical experimental reagents. SL, CZ and JW conducted bioinformatics analysis on RNA-seq and ChIP-seq data. JW, JF and JZ designed experiments. JZ provided the conceptual idea and overall supervision of the project.

## Figure Legends

**Supplemental Figure 1 Legend.**
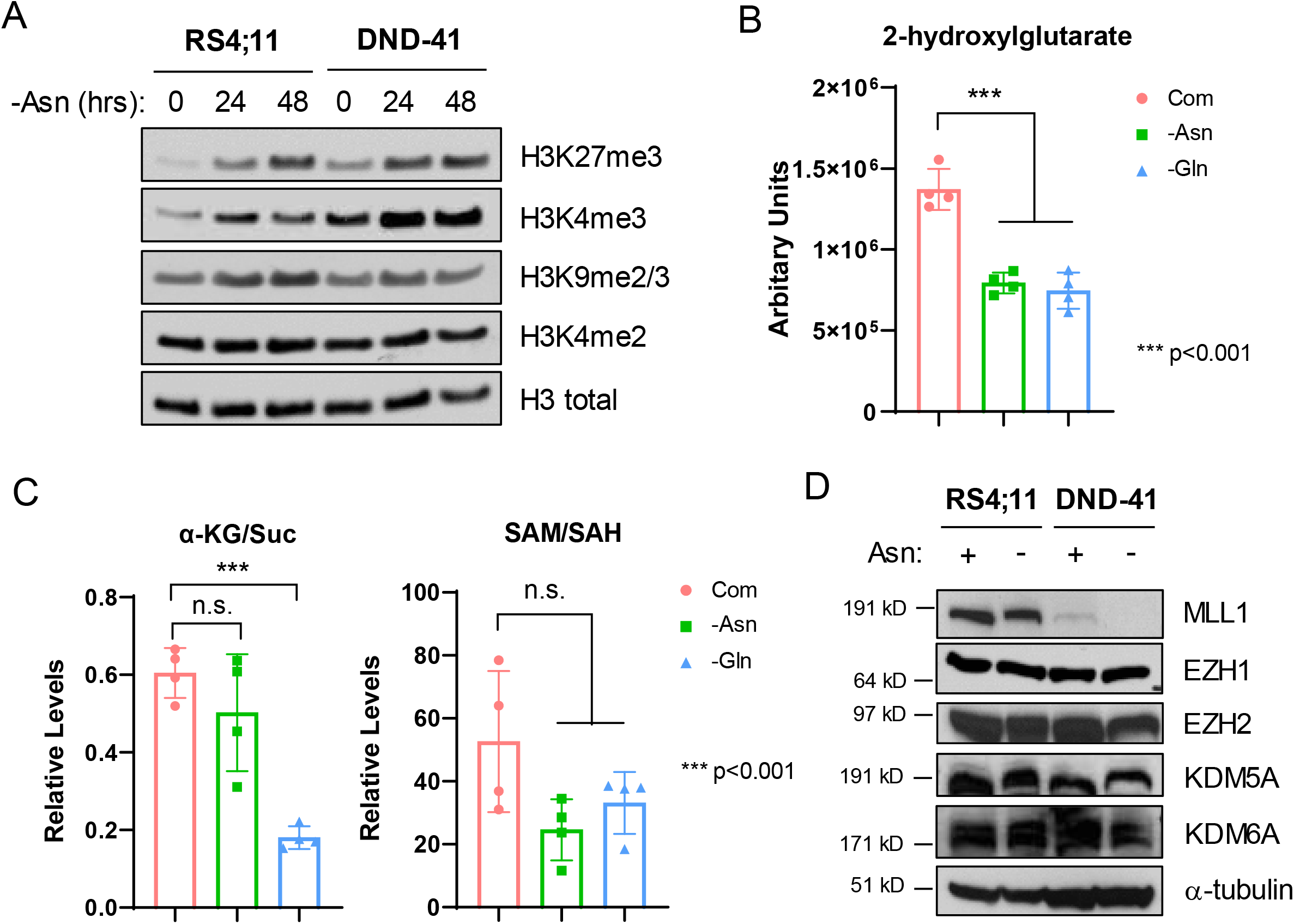
A. RS4;11 and DND-41 cells were subjected for asparagine starvation. Nuclear extracts were harvested at 24 and 48 hours, and were subjected to Western blotting for H3K27me3, H3K4me3, H3K9me2/3, H3K4me2 and total H3. B. RS4;11 cells were subjected to asparagine or glutamine starvation for 24 hours. Intracellular 2-hydroxylglutarate levels were determined by LC-MS (n=4). C. RS4;11 cells were subjected to asparagine or glutamine starvation for 24 hours. Intracellular α-ketoglutarate (α-KG), succinate (Suc), SAM and SAH levels were determined by LC-MS. The ratio of α-KG/Suc and SAM/SAH was calculated based on arbitrary units (n=4). D. RS4;11 and DND-41 cells were subjected to asparagine starvation for 24 hours. Protein extracts were subjected to Western blotting for MLL1, EZH1, EZH2, KDM5A, KDM6A and α-tubulin. Results in panel B and C were shown as mean ± SD (standard derivation). p values were determined by using Student’s two-tailed unpaired t-test.

**Supplemental Figure 2 Legend.**
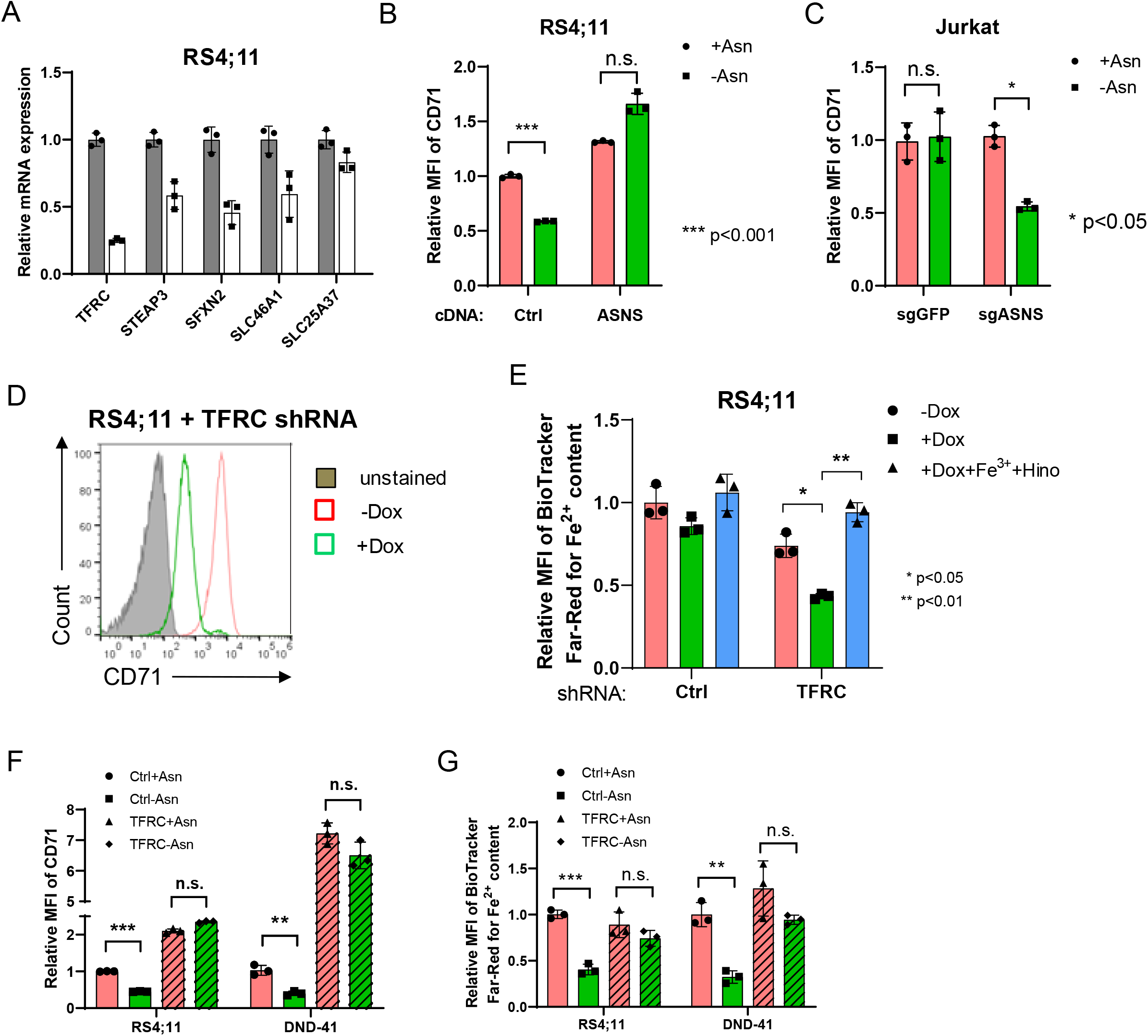
A. RS4;11 cells were subjected to asparagine starvation for 24 hours. RNA was harvested and subjected to qPCR analysis for *TFRC, STEAP3, SFXN2, SLC46A1 and SLC25A37*. B. RS4;11 cells expressing an empty vector (Ctrl) or *ASNS* cDNA were subjected to asparagine starvation for 24 hours. Cell surface expression of TFRC/CD71 was determined by flow cytometry and the mean fluorescence intensity (MFI) was quantified. C. Jurkat cells expressing a control guide RNA (sgGFP) or a guide RNA for human *ASNS* (sgASNS) were subjected to asparagine starvation for 24 hours. Cell surface expression of TFRC/CD71 was determined by flow cytometry and the mean fluorescence intensity (MFI) was quantified. D. RS4;11 cells expressing a doxycycline-inducible shRNA for *TFRC* gene were treated with doxycycline (1 µg/mL) for 48 hours. Cell surface expression of TFRC/CD71 was determine by flow cytometry analysis. E. RS4;11 cells expressing a doxycycline-inducible control shRNA or *TFRC* shRNA were treated with doxycycline (1 µg/mL) for 48 hours with or without Fe^3+^ supplementation (10 µM) in the presence of Hinokitiol (0.75 µM). Intracellular Fe^2+^ content was determined by BioTracker Far-Red staining and flow cytometry analysis. E. RS4;11 and DND-41 cells expressing an empty vector (Ctrl) or *TFRC* cDNA were subjected to asparagine starvation for 24 hours. Cell surface expression of TFRC/CD71 was determined by flow cytometry analysis. G. RS4;11 and DND-41 cells expressing an empty vector (Ctrl) or *TFRC* cDNA were subjected to asparagine starvation for 24 hours. Intracellular Fe^2+^ content was determined by BioTracker Far-Red staining and flow cytometry analysis. Results in panel B, C, E, F and G were shown as mean ± SD (standard derivation). p values were determined by using Student’s two-tailed unpaired t-test.

**Supplemental Figure 3 Legend.**
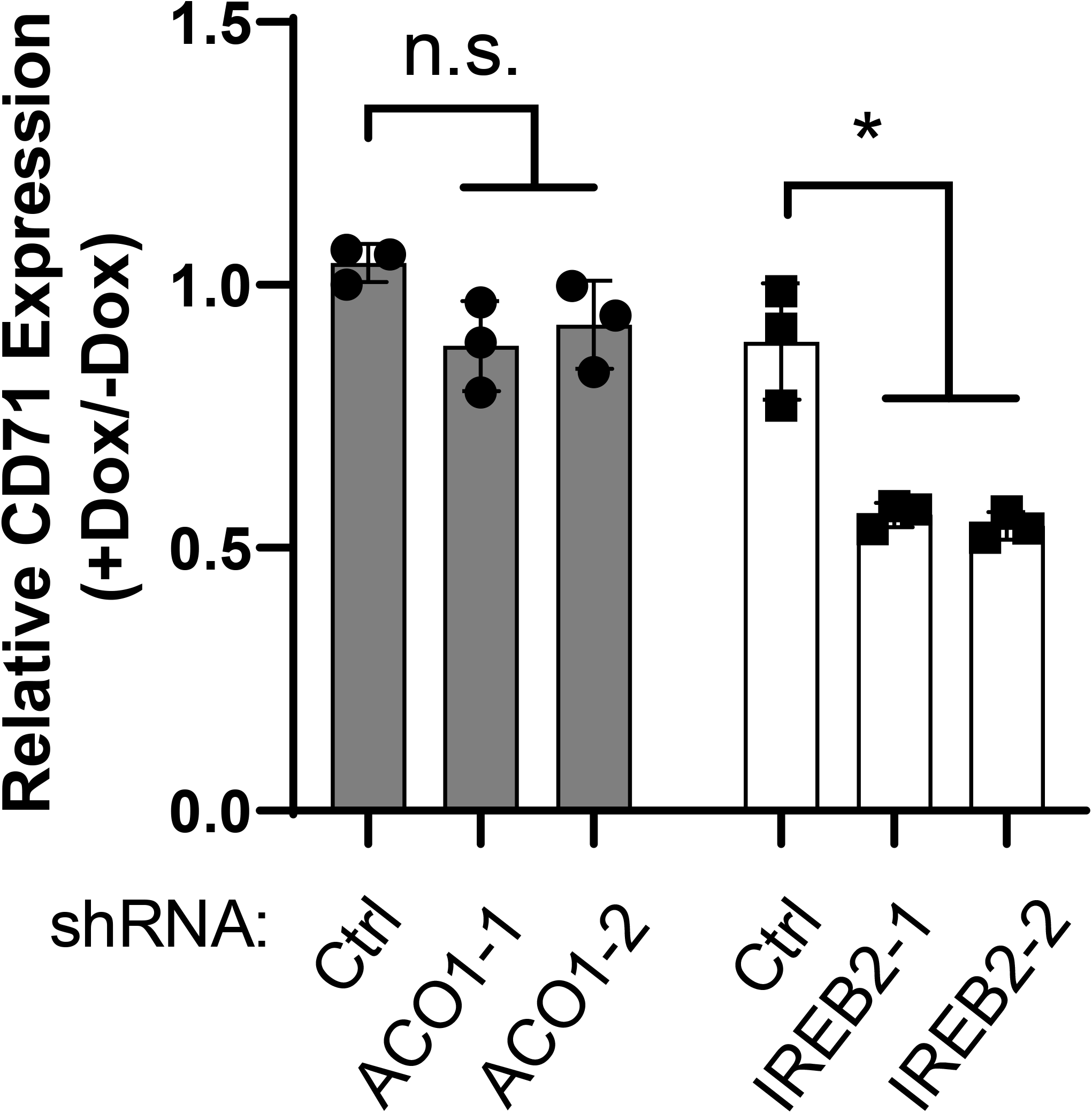
RS4;11 cells expressing doxycycline-inducible control, *ACO1* or *IREB2* shRNA were treated with or without doxycycline (1 µg/mL) for 48 hours. The mRNA expression of TFRC/CD71 was determined by qPCR analysis. For *ACO1* and *IREB2* genes, two independent hairpins were chosen. Results were shown as mean ± SD (standard derivation). p values were determined by using Student’s two-tailed unpaired t-test.

**Supplemental Figure 4 Legend.**
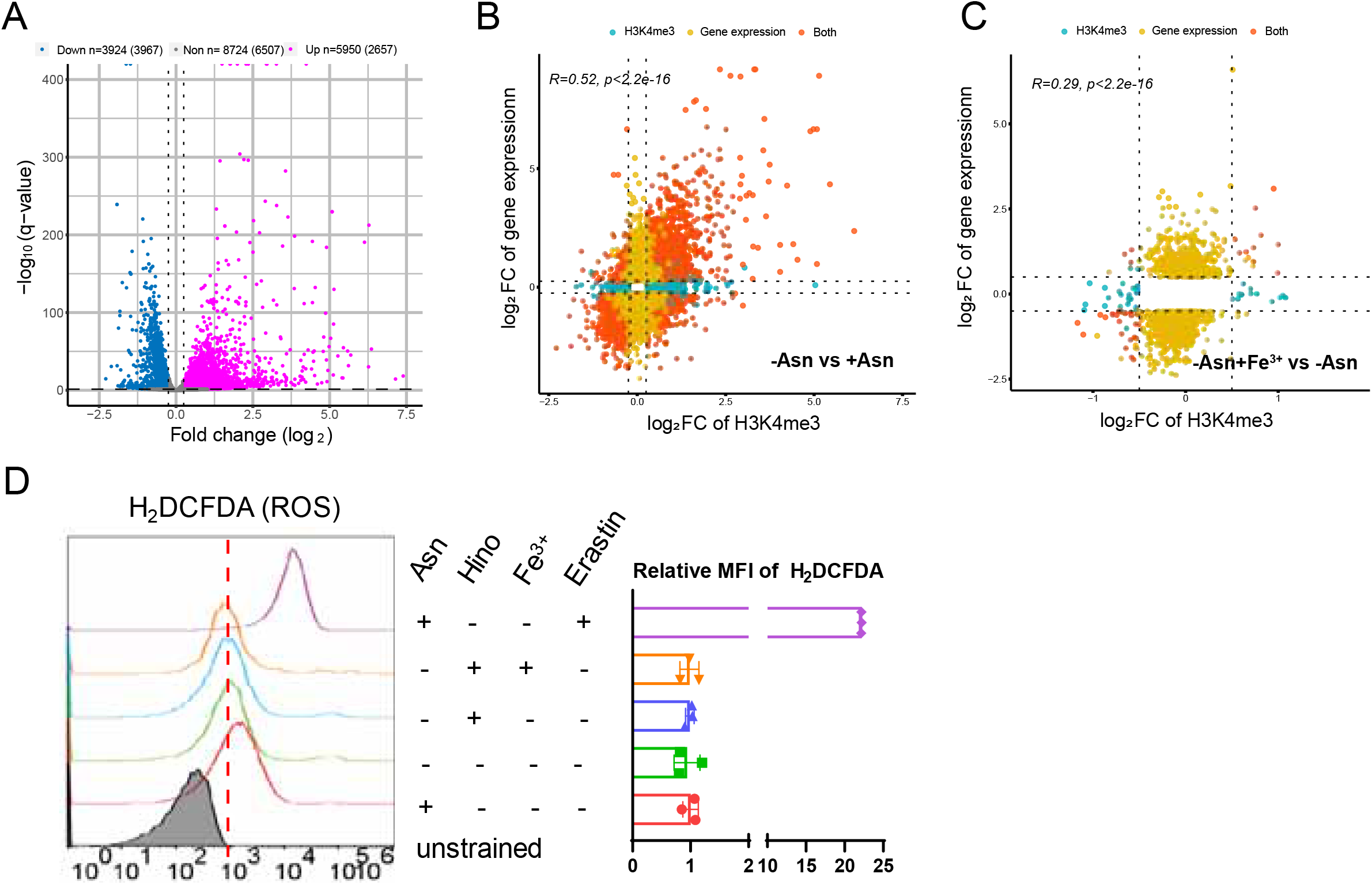
A. RS;41 cells were cultured in asparagine-replete or -deficient media for 24 hours. H3K4me3 ChIP-seq analysis were performed. Volcano plot of differential expression peaks (FDR<0.05, |log_2_FC|>0.25) were illustrated. The number of peaks and corresponding genes was listed on the top of the plot. B. RS4;11 cells were cultured in the presence of asparagine (+Asn), in the absence of asparagine (-Asn) or in the absence of asparagine but with Fe^3+^ supplementation (-Asn+Fe^3+^) condition for 24 hours. H3K4me3 ChIP-seq and whole cell RNA-seq were performed. Scatter plot of differential expression peaks and differential expression genes in the −Asn vs +Asn condition was shown. C. Scatter plot of differential expression peaks and differential expression genes in the −Asn+Fe^3+^ vs −Asn condition was shown. D. RS4;11 cells were subjected to different treatments indicated in the legend for 24 hours. Intracellular ROS was determined by H_2_DCFDA staining and flow cytometry analysis. Erastin (10 µM) was used as a positive control for H_2_DCFDA staining. Fe^3+^ (10 µM); Hinokitiol (0.75 µM). Results on the right panel were shown as mean ± SD (standard derivation).

